# Pulvinar Slits: Cellulose-deficient and De-Methyl-Esterified Homogalacturonan-Rich Structures in a Legume Motor Cell

**DOI:** 10.1101/2022.03.10.483846

**Authors:** Masahiro Takahara, Satoru Tsugawa, Shingo Sakamoto, Taku Demura, Miyuki T Nakata

## Abstract

The cortical motor cells (CMCs) in a legume pulvinus execute the reversible deformation in leaf movement that is driven by changes in turgor pressure. In contrast to the underlying osmotic regulation proper, the cell wall structure of CMCs that contributes to the movement has yet to be characterized in detail. Here, we report that the cell wall of CMCs has circumferential slits with low levels of cellulose deposition. This structure is unique and distinct from that of any other primary cell walls reported so far; thus, we named them “pulvinar slits.” Notably, we predominantly detected de-methyl-esterified homogalacturonan inside pulvinar slits, with a low deposition of highly methyl-esterified homogalacturonan, as with cellulose. In addition, Fourier-transform infrared spectroscopy analysis indicated that the cell wall composition of pulvini is different from that of other axial organs, such as petioles or stems. Moreover, monosaccharide analysis showed that pulvini are pectin-rich organs like developing stems and that the amount of galacturonic acid in pulvini is greater than in developing stems. Computer modeling suggested that pulvinar slits facilitate anisotropic extension in the direction perpendicular to the slits in the presence of turgor pressure. When tissue slices of CMCs were transferred to different extracellular osmotic conditions, pulvinar slits altered their opening width, indicating their deformability. In this study, we thus characterized a distinctive cell wall structure of CMCs, adding new knowledge to repetitive and reversible organ deformation as well as the structural diversity and function of the plant cell wall.

## Introduction

A legume pulvinus is a joint-like thickening at the base of a petiole, a petiolule, or a leaflet that plays a central role in leaf movement. Pulvinus-actuated leaf movements include circadian movements completed over a scale of hours (Darwin and Darwin, 1880), blue light–induced movements within minutes (Koller, 1990), and touch-stimulated movements of Mimosoideae within seconds (Weintraub, 1952). During movement, a pulvinus is repeatedly and reversibly deformed. These repetitive and reversible steps are a unique feature that distinguish pulvinus-based movements from other types of organ movement driven by irreversible growth or by mechanical instability (Dumais and Forterre, 2012; Geitmann, 2016).

Since the cellular membrane acts as a semi-permeable barrier, a change in cell volume is the result of water flux between the inside and the outside of the cell (and the vacuole) and is triggered by the difference in water potential between the two sides (Morris and Blyth, 2019; Nakata and Takahara, 2022). In inner tissues and in a microenvironment, water potential almost equals osmotic potential; cells swell when exposed to a hypotonic condition, while they contract under a hypertonic condition. Plant cells regulate their osmotic potential in a variety of ways, for example, via the gate opening or closure of ion channels or by increasing or decreasing the concentration of osmotic compounds through metabolic pathways. Importantly, excessive expansion is blocked by the physical scaffold provided by the cell wall. When the cell volume increases in a hypotonic solution, cell walls act as a load-bearing structure. The difference between the osmotic potential inside and outside the cell triggers cell expansion and generation of in-plane tensile force in the cell wall. When the in-plane tensile force is balanced with the load-bearing strength of the cell wall, the water potential inside the cell equals that outside the cell to stop the water flux. In this situation, the difference between the osmotic potential inside and outside the cell corresponds to the “turgor pressure” of the cell.

The primary cell wall (PCW) consists mainly of cellulose, xyloglucan, and pectin. Among PCW components, multiple cellulose polymers are bundled into microscale fibers, namely, cellulose microfibrils, to provide tensile stiffness and strength to the PCW. In axially growing cells, the orientation of cellulose microfibrils often exhibits a bias perpendicular to the direction of expansion. In addition, xyloglucan and pectin contribute to cell wall extensibility by interacting with cellulose microfibrils (Cosgrove, 2005; Smithers et al., 2019). In growing cells, cell expansion is triggered by loosening of the cell wall by changing connections between cellulose microfibrils (Cosgrove, 2005). In the context of reversible deformation, the connections between cellulose microfibrils are thought to not be altered and to behave as an elastic material, with changes in osmotic potential triggering cell expansion or contraction (Nakata and Takahara, 2022). The difference of mechanical behavior suggests that cell wall properties of reversibly deforming cells, like motor cells, differ from those of growing cells.

Pulvinus-driven movements are independent from growth (Darwin and Darwin, 1880). The pulvinus comprises several mature tissues, including the epidermis, cortical motor cells (CMCs), and vascular tissues (Fleurat-Lessard and Bonnenmain, 1978; Fleurat-Lessard and Roblin, 1982; Moysset and Simon, 1991). CMCs have a central role in the reversible deformation of the pulvinus. The CMCs that shrink when the pulvinus flexes are called extensor motor cells, while CMCs on the opposite side are called flexor cells. In the early 1900s, changes in turgor of CMCs were proposed to be critical for leaf movements (Weintraub, 1952). Later, the bulk moduli of extensibility estimated in a study using runner bean (*Phaseolus coccineus*) indicated that the difference in osmotic concentration between the inside and the outside of a CMC is a major determinant of how much the CMC volume will change and how much leaf movement will be actuated (Mayer et al., 1985). Therefore, the changes in osmotic concentrations in CMCs have received much attention in various legume species with pulvini (reviewed in Moran, 2007).

Mayer et al. also reported anisotropic extension to the proximo-distal (PD) direction in an intact pulvinus during nyctinastic movement and in slices of CMCs exposed to different concentrations of mannitol (Mayer et al., 1985). The extension anisotropy of CMCs was confirmed by recent studies of other species (Koller and Zamski, 2008; Sleboda et al., 2022). According to these findings, the cell wall of CMCs is thought of as having physical properties suitable for significant changes in volume with a bias in the PD direction. It has been reported that cellulose microfibrils are aligned in a circumferential orientation along the adaxial/abaxial (AdAb)-mediolateral (ML) plane in the cell wall of CMCs in *P. coccineus* and sensitive plant (*Mimosa pudica*) (Mayer et al., 1985; Sleboda et al., 2022). Indeed, the cell wall exhibits great elasticity in the direction along cellulose microfibrils (more than 1 GPa) compared to the range of isotropic turgor pressure in a given tissue (approximately 1 MPa), which acts as a mechanical constraint for cell elongation, resulting in the extension anisotropy of CMCs. However, the anisotropy of cellulose microfibrils is not sufficient to explain the observed differences between the cell wall of CMCs and that of growing cells.

In this study, to better understand the mechanical basis of reversible deformation, we investigated the cell wall structure of CMCs in detail. Since the shape of CMCs is a polyhedron close to a sphere, we produced slices cut from a pulvinus by hand-sectioning or agarose sectioning to observe a large area of the cell wall and visualize the cellulose localization. We also acquired optical sections with a confocal laser-scanning microscope. Surprisingly, we discovered that the cell wall of CMCs is not homogeneous but has previously undescribed slit-like structures with less cellulose that we designated “pulvinar slits.” We also determined that pulvinar slits are widely conserved among legume species. Inside pulvinar slits, we detected the abundant deposition of de-methyl-esterified homogalacturonan (HG), while highly methyl-esterified HG was not detected. Furthermore, we show that the pulvinus is a pectin-rich organ in comparison to other axial organs and confirmed that the PD extensibility of the pulvinus is higher than that of other axial organs by a mechanical test in four legume species. Computational simulations suggested that the presence of pulvinar slits causes anisotropic extension biased in the PD direction without any other mechanical anisotropy like the biased orientation of cellulose microfibrils. We also determined that changes in the opening width of pulvinar slits accompany changes in cell volume induced by exchanging external solutions (from hypotonic to hypertonic or vice versa). The discovery of pulvinar slits in this study sheds light on a new mechanical model for repetitive and reversible organ deformation.

## Results

### CMCs have circumferential slits with less cellulose

To identify the cell wall structure of CMCs, we stained pulvini with calcofluor white, which stains cell wall components such as cellulose, in vertical and cross sections using pulvini freshly harvested from panicledleaf ticktrefoil (*Desmodium paniculatum*), belonging to the Papilionoideae family (Figure 1A–F). Notably, in CMC vertical sections, we observed several slits with no significant fluorescence when stained by calcofluor white that run in a circumferential orientation in the adaxial/abaxial (AdAb)-mediolateral (ML) plane (Figure 1C, G). In vertical sections, we noticed small spots with no staining but no slits (Figure 1D, G). To determine whether circumferential slits are specific to CMCs, we repeated the staining protocol on cortical cells collected from the boundary between a pulvinus and a petiole and from a vertical section of a petiole (Figure 1E, F). In both cases, we visualized small spots lacking staining but did not see circumferential slits (Figure 1E, F). Since we only observed these circumferential slits in pulvinus CMCs, we designated the circumferential slits as pulvinar slits.

**Figure 1.**
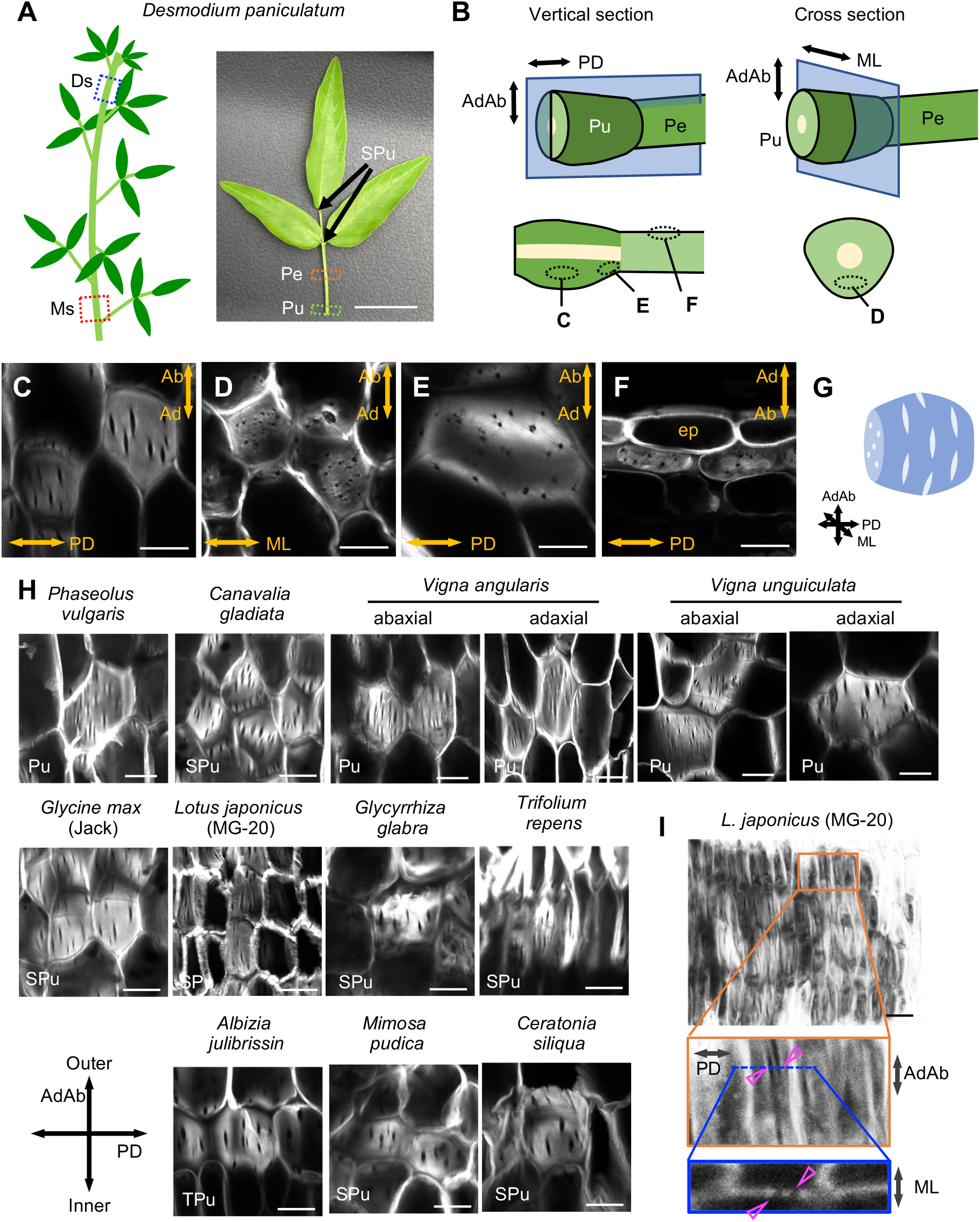
Calcofluor white staining of pulvinar cortical motor cells (CMCs) in various legume species. (A) Schematic diagram (left) and photograph (right) of a *Desmodium paniculatum* leaf. Pu, primary pulvinus; SPu, secondary pulvinus; Pe, petiole; Ds, developing stem; Ms, mature stem. (B) Schematic diagrams of the collected pulvini and petiole samples for microscopy. Letters C to F indicate the positions visualized in the corresponding panels. (C-F) Sections of calcofluor white–stained *D. paniculatum* pulvini and petioles by confocal laser-scanning microscopy. AdAb, adaxial-abaxial axis; PD, proximo-distal axis; ML, medial-lateral axis; ep in F indicates epidermis. (G) A model of pulvinar slits and small spots in the cell wall of a CMC. TPu, a tertiary pulvinus. (H) Vertical sections of calcofluor white–stained pulvinar CMCs from 11 legume species. The vertical direction of all images corresponds to the adaxial-abaxial axis. The top views correspond to the outside of a pulvinus, and the bottom corresponds to the inside. (I) Whole-mount projection of calcofluor white-stained *L. japonicus* CMCs (top). Middle, magnified view of the region highlighted by the orange square. Bottom, optical Z-section shown as the blue line in the middle panel. Arrowheads indicate pulvinar slits. Scale bars, 2 cm (A), or 20 μm (C-F, H, I).

To investigate the extent of conservation of pulvinar slits among legumes, we stained vertical sections of pulvini from various legume species with calcofluor white. The Fabaceae family comprises two major subfamilies: the Papilionoideae and the Caesalpinioideae (Azani et al., 2017). Importantly, we detected slit structures in pulvinar CMCs from all 11 species selected and tested from these two subfamilies (Supplemental Table S1, Figure 1H). In addition, we observed pulvinar slits in both adaxial and abaxial pulvinar CMCs (Figure 1H). To make sure that the slits were not artifacts arising during sectioning, we performed whole-mount staining. Again, we clearly observed slits in CMCs from whole-mount-stained samples, with the slits located at the cell periphery (Figure 1I). We therefore concluded that pulvinar slits are an innate cell wall structure of CMCs.

To validate the observation that a pulvinar slit is a region with lower cellulose deposition than its surrounding region, we used cellulose-specific dyes, namely, pontamine and immunolabeling with an antibody against type A carbohydrate-binding module 3 (CBM3a). With pontamine staining, we detected circumferential slits similar to those seen with calcofluor white staining in pulvinar CMCs (Figure 2A). Immunolabeling with CBM3a predominantly stains crystalline cellulose as speckles, but we also found slit-like structures with lower signals (Figure 2B). Double staining with CBM3a and calcofluor white indicated that slits lacking the CBM3a signal colocalize with pulvinar slits, the structures unstained by calcofluor white (Figure 2B, arrowheads). These observations indicated that pulvinar slits are regions characterized by an extremely low cellulose deposition.

**Figure 2.**
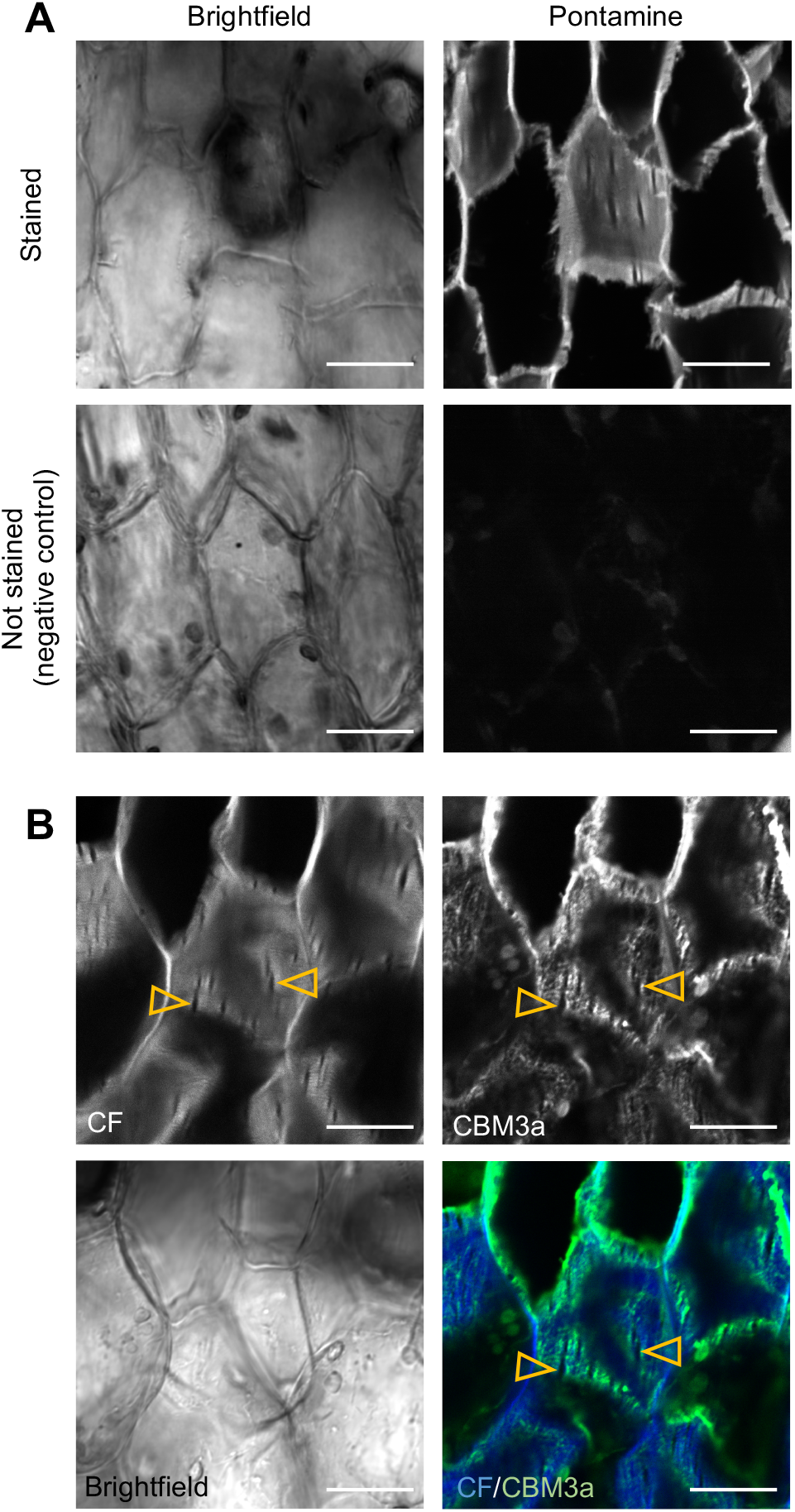
Visualization of cellulose in *D. paniculatum* pulvinar CMCs. (A) Pontamine-stained CMCs of a *D. paniculatum* pulvinus (vertical section). (B) Co-localization of calcofluor white signal (CF) and staining with the CBM3a antibody (recognizing crystalline cellulose) in CMCs in a vertical section of a *D. paniculatum* pulvinus. Arrowheads, pulvinar slits. Scale bars, 20 μm.

Taken together, we conclude that pulvinar slits are a structure specifically formed on the PCW of both extensor and flexor (adaxial and abaxial) CMCs, with less cellulose than the surrounding cell wall, and that they are widely conserved among legume species.

### De-methyl esterification of homogalacturonan is associated with pulvinar slits

To further characterize pulvinar slits, we examined the spatial pattern of other major PCW components by immunohistological analysis. HG is the most abundant pectin, while xyloglucan is the main hemicellulose detected in the PCW. HG is methyl-esterified during its biosynthesis and later de-methyl-esterified during maturation. We detected a strong signal with the LM19 antibody, which labels de-methyl-esterified HG, inside pulvinar slits with a stronger signal close to the boundary with the rest of the PCW (Figure 3A, B). By contrast, the LM20 antibody, which labels highly methyl-esterified HG, showed no signal in pulvinar slits (Figure 3A, C). We also labeled XXXG-type xyloglucan with the LM15 antibody, which revealed dispersed signals throughout CMCs (Figure 3A). We detected a faint signal with the 2F4 antibody, which labels calcium-crosslinked de-methyl-esterified HG, although the signal appeared to be excluded from pulvinar slits (Supplemental Figure S1A). We did not observe a signal when sections were incubated only with the secondary antibody (Supplemental Figure S1B). These results thus suggested that de-methyl esterification of homogalacturonan, probably without calcium crosslinking, is associated with pulvinar slits.

**Figure 3.**
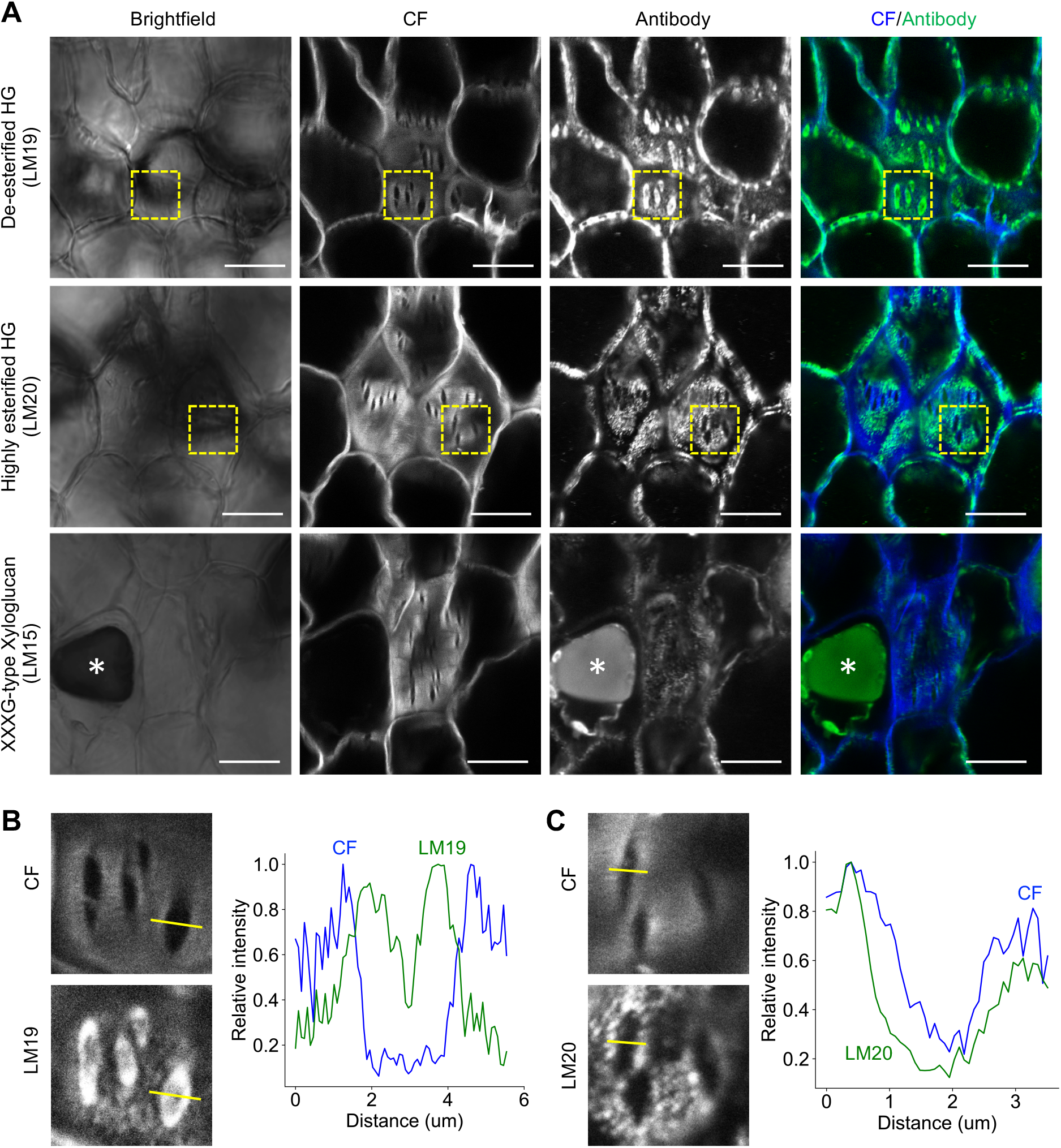
Immunohistological analyses of *D. paniculatum* pulvinar CMCs. (A) Immunolabeled CMCs with antibodies against primary cell wall (PCW) components in vertical sections. These samples were co-stained with calcofluor white (CF). Asterisk indicates a tannin vacuole. (B-C) Magnified views of dashed boxes in (A) (left) and profiles of relative intensities for calcofluor white (blue) and antibody signal (green) along the yellow lines in the magnified images (right). Scale bars, 20 μm (A).

To better assess the composition of the pulvinus cell wall, we used Fourier-transform infrared spectroscopy (FTIR) and monosaccharide analysis to compare the cell wall from pulvini to that of other axial organs. Using the absorbance profile from FTIR, we identified several peaks (e.g., 778 and 1317 cm^−1^) and a range of 1620–1550 cm^−1^ that are specific to pulvini (Figure 4A). A principal component analysis revealed that the cell wall properties obtained from FTIR can be classified into three groups (shown as the dashed circles in Figure 4B), with pulvinar organs forming a group distinct from all other axial organs (Figure 4B). The monosaccharide composition of pulvinar organs (primary pulvinus and secondary pulvinus) was also analyzed in detail. The xylose content mainly related to xylan in the secondary cell wall of pulvini and developing stems was lower than that of petioles and mature stems (Figure 4C, Table 1). By contrast, pectin-related monosaccharides, such as galactose, arabinose, rhamnose, and galacturonic acid, were more abundant in pulvini than in petioles and in mature stems (Figure 4C, Table 1). Galactose, arabinose, and rhamnose were also abundant in developing stems, whereas the abundance of galacturonic acid of pulvini was higher than that of developing stems (Figure 4C). Among pectins, HG is composed of only galacturonic acid, but rhamnogalacturonan-I (RG-I) and RG-II are composed of other monosaccharides (e.g., galactose, arabinose, and rhamnose) in addition to galacturonic acid. Therefore, these results suggest that, despite being mature organs, the cell wall composition of pulvini is relatively similar to that of developing stems rather than that of mature organs (petioles and mature stems) and is rich in pectin (probably HG).

**Figure 4.**
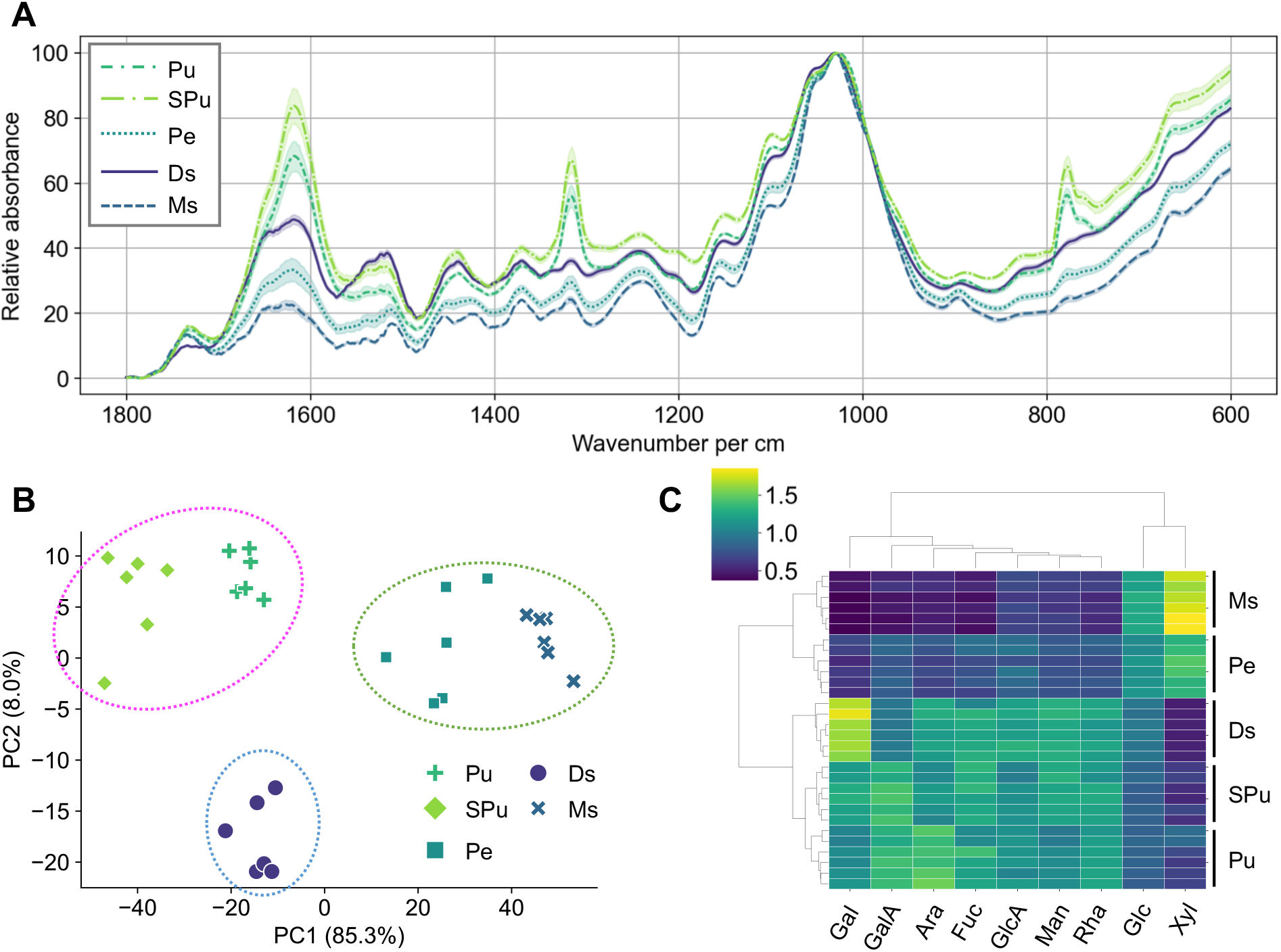
Cell wall assay of *D. paniculatum* axial organs. (A) FTIR spectra of cell wall components obtained from five different axial organs: primary pulvini (Pu), secondary pulvini (SPu), petioles (Pe), developing stems (Ds), and mature stems (Ms), as six biological replicates. Data are shown as the mean and 95% confidence interval. (B) Principal component analysis applied to the FTIR spectra shown in A. (C) Heatmap representation with corresponding dendrogram of the relative amount of each monosaccharide in each organ. Color indicates the ratio of the mean for each monosaccharide. Gal, galactose; GalA, galacturonic acid; Ara, arabinose; Fuc, fructose; GlcA, glucuronic acid; Man, mannose; Rha, rhamnose; Glc, glucose; Xyl, xylose.

**Table 1.** The amount of cell-wall components (μg/gAIR) in aerial axial organs

|  | Primary pulvinus |  | Secondary pulvinus |  | Petiole |  | Developing stem |  | Mature stem |  |
| --- | --- | --- | --- | --- | --- | --- | --- | --- | --- | --- |
|  | Mean | SD | Mean | SD | Mean | SD | Mean | SD | Mean | SD |
| Glc*** | 228.97 | 8.27<sup>ab</sup> | 180.03 | 5.46<sup>a</sup> | 345.59 | 20.16<sup>bc</sup> | 221.73 | 15.08<sup>ab</sup> | 401.62 | 19.84<sup>c</sup> |
| GalA*** | 157.30 | 5.98<sup>a</sup> | 126.41 | 8.79<sup>ab</sup> | 109.45 | 6.01<sup>abc</sup> | 96.89 | 4.89<sup>bc</sup> | 66.94 | 10.33<sup>c</sup> |
| Xyl*** | 51.73 | 9.12<sup>ab</sup> | 33.49 | 3.43<sup>a</sup> | 108.06 | 8.60<sup>b</sup> | 30.05 | 1.65<sup>a</sup> | 143.79 | 10.76<sup>b</sup> |
| Gal*** | 46.61 | 1.73<sup>ab</sup> | 41.87 | 3.17<sup>abc</sup> | 30.51 | 3.02<sup>bc</sup> | 61.40 | 1.66<sup>a</sup> | 20.64 | 2.59<sup>c</sup> |
| Man*** | 41.92 | 1.97<sup>ab</sup> | 39.57 | 1.49<sup>abc</sup> | 34.40 | 1.73<sup>bc</sup> | 43.92 | 1.06<sup>a</sup> | 30.57 | 2.76<sup>c</sup> |
| Ara*** | 38.03 | 1.67<sup>a</sup> | 23.29 | 2.09<sup>ab</sup> | 21.58 | 1.82<sup>bc</sup> | 27.84 | 0.76<sup>ac</sup> | 16.25 | 1.77<sup>b</sup> |
| Rha*** | 14.08 | 0.34<sup>a</sup> | 11.17 | 0.92<sup>ab</sup> | 10.25 | 0.67<sup>bc</sup> | 12.03 | 0.14<sup>ac</sup> | 8.13 | 0.99<sup>b</sup> |
| GlcA*** | 2.86 | 0.18<sup>a</sup> | 2.17 | 0.18<sup>ab</sup> | 2.51 | 0.28<sup>ab</sup> | 2.67 | 0.12<sup>a</sup> | 2.04 | 0.17<sup>b</sup> |
| Fuc*** | 2.88 | 0.29<sup>a</sup> | 2.41 | 0.12<sup>ab</sup> | 2.14 | 0.26<sup>bc</sup> | 2.52 | 0.23<sup>ac</sup> | 1.33 | 0.15<sup>b</sup> |
| Total*** | 584.37 | 15.82<sup>ab</sup> | 460.40 | 17.74<sup>a</sup> | 664.48 | 30.48<sup>b</sup> | 499.05 | 23.41<sup>a</sup> | 691.31 | 36.70<sup>b</sup> |

n = 6; \*\*\*p < 0.001 by the Kruskal-Wallis H-test; a-c, different characters indicate significantly different (p<0.05) by pairwise test for multiple comparisons of mean rank sums (Dunn's test; Holm's method)

Since the composition of the pulvinar cell wall was clearly different from that of other axial organs, we wondered if pulvini might exhibit mechanical differences from other axial organs. To test this hypothesis, we analyzed the PD extensibility under compression using the principle of the Poisson’s effect, in which a material generally expands in directions perpendicular to the direction of compression (Supplemental Figure S2). Specifically, because pulvini are thought to be analogous to petiole-like organs, petiolules, based on the *petiolule-like pulvinus* mutant phenotype in barrel clover (*Medicago truncatula*) (Zhou et al., 2012), we focused on the difference between pulvini and petioles. In *D. paniculatum*, we detected load oscillations, probably due to tearing of the epidermis and separation of vascular bundles from CMCs, in pulvini (Supplemental Figure S3A, S4). Similar oscillations were detected in developing stems, but not in petioles and rarely in mature stems (Supplemental Figure S3A, S4). The length of pulvini along the PD axis increased slowly until the strain reached 0.3 but extended much more for strain values above 0.4 (Figure 5A–B). The length of developing stems increased, but not as much as that of pulvini (Figure 5A). We did not observe a statistically significant extension of petioles, while mature stems only extended slightly under greater strain (Figure 5A). For a strain value of 0.5, the extension rate of pulvini was higher than that of other axial organs (Figure 5C). Similarly, we determined that pulvini display substantial PD extension, which was accompanied by load oscillation in three other legume species (Figure 5D, Supplemental Figure S3B–C). These results indicate that the pulvinus displays excellent PD extensibility at the organ level.

**Figure 5.**
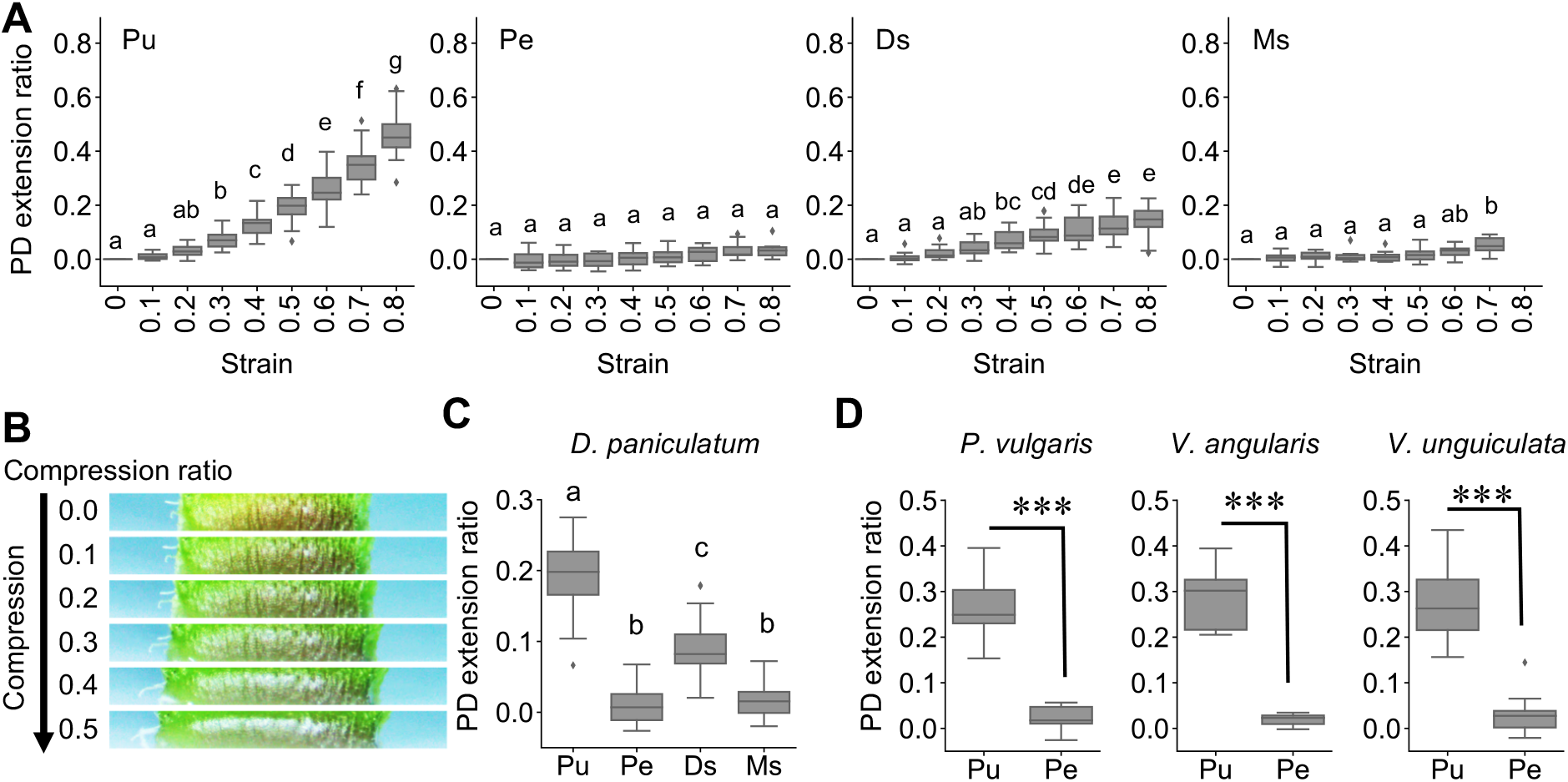
Compression test of pulvini and other axial organs. (A) Ratio of the PD extension of the axial organs for each strain (compression ratio) applied. n = 18 (pulvini, Pu), 10 (petioles, Pe), 15 (developing stems, Ds), 10 (mature stems, Ms, other than strain = 0.7), and 8 (mature stems, strain = 0.7). Different letters indicate significant differences (Tukey’s honestly significant difference [HSD] test, family-wise error rate = 0.05). (B) Serial images of side views of *D. paniculatum* pulvini during compression with a creep meter. (C) Ratio of the PD extension of the axial organs in *D. paniculatum* when the applied strain is 0.5. (D) Ratio of the PD extension of axial organs in three other species when the applied strain is 0.5. n = 15 (*P. vulgaris* petioles), 16 (*P. vulgaris* pulvini), 12 (*V. angularis*), and 11 (*V. unguiculata*). \*\*\**p* < 0.001 by two-sided Welch’s *t*-test. The calculated t-statistics for *P. vulgaris, V. angularis*, and *V. unguiculata* are 12.20, 9.04, and 9.05, respectively.

### Pulvinar slits in the cell wall have a potential effect on the PD extensibility of pulvini

To explore the mechanical role of pulvinar slits in the turgor-triggered morphological changes of CMCs, we implemented the finite-element method (FEM) to determine how cell shape is affected by the slits under turgor pressure. A previous study using a single cell layer sheet of onion (*Allium cepa*) peels indicated that cellulose mainly contributes to in-plane rigidity and plasticity, while pectin contributed very little (Zhang et al., 2019). Therefore, we hypothesized that the region near pulvinar slits would be mechanically weak during deformation of CMCs under turgor. We modeled the cross-sectional shape of CMC in a PD-AdAb plane or PD-ML plane as shapes similar to a hexagon with several elliptical holes (Figure 6A). To set an accurate slit size for simulations, we measured cell length and the length and width of the slits in 59 randomly selected slits from eight species, yielding slit lengths ranging from 2.03 to 10.06 μm and a slit width ranging from 0.32 to 2.26 μm. With a cell length of 20.6–46.1 μm, slit length was between 7.2% and 25.4% of the overall cell length, with a mean of about 15%, while slit width was between 0.9% and 9.9% of cell length, with a mean of 3% (Figure 6B). We also calculated the slit index (length/width), which was 2.46–16.60 with a mean of about 6.14.

**Figure 6.**
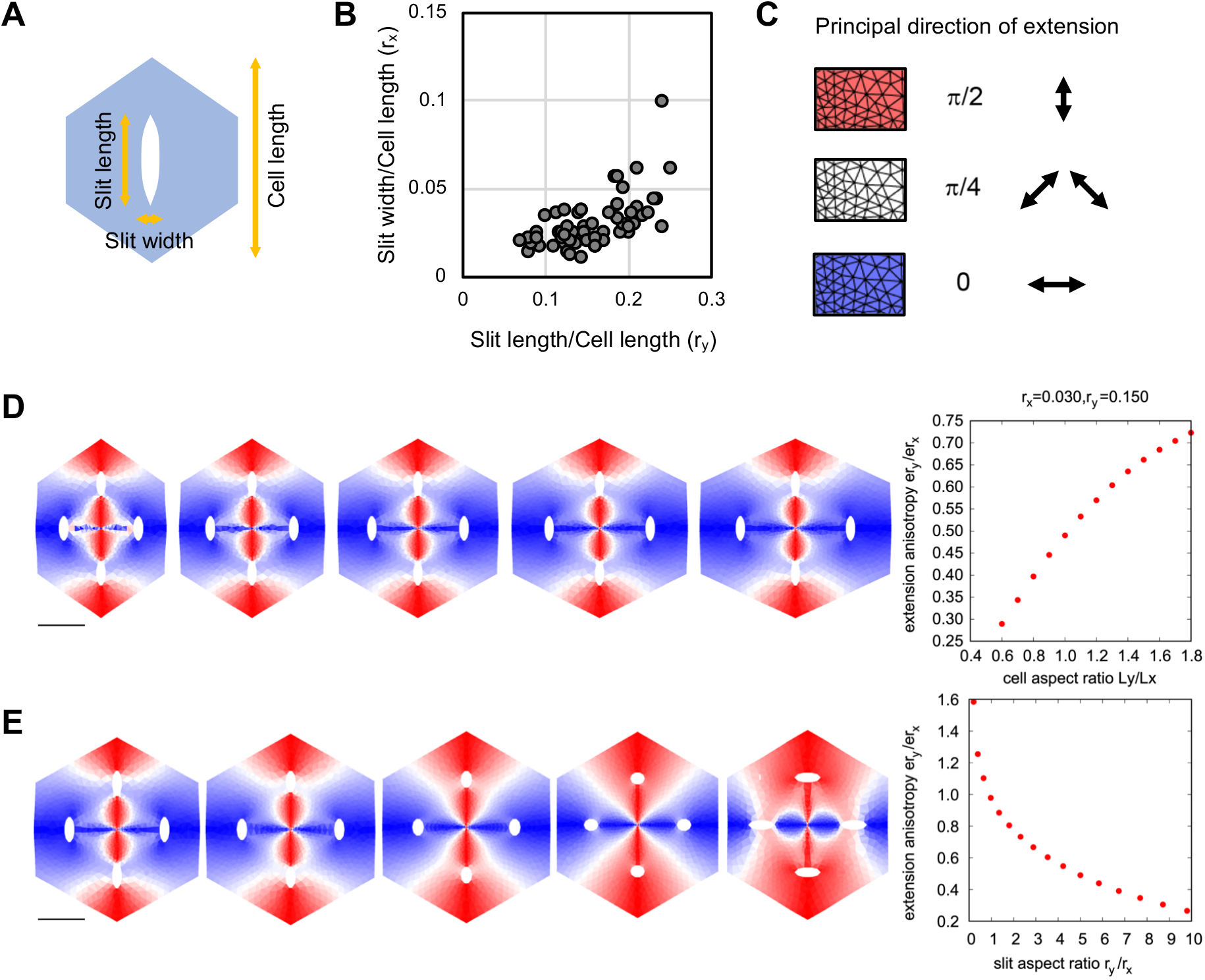
Computational modeling of the expansion of a hexagonal cell with slits. (A) Schematic diagram of the hexagonal cells with definition of terms used. (B) Relative length and width of 59 slits from eight species: *D. paniculatum, P. vulgaris, V. angularis, V. unguiculata, G. max* (Jack), *A. julibrissin, M. pudica*, and *C. siliqua*. (C) Color key for principal direction of growth shown in D and E. 0, horizontal plane; π/2, vertical plane. (D-E) Results of FEM simulations to clarify the effect of x-y aspect ratio of cells (D) and slit direction (E). Extension anisotropy was calculated from the extension ratio along the horizontal (x) axis (er_x_) and along the vertical (y) axis (er_y_). The cell aspect ratio was calculated from cell length along the x-axis (L_x_) and along the y-axis (L_y_). Slit aspect ratio was calculated from slit length along the x-axis (r_x_) and along the y-axis (r_y_).

Based on these shape parameters in experiments as a standard slit property, we generated several cell models with different cell shapes and with different slit properties (Figure 6C–E, Supplemental Figure S5). Using FEM, we simulated cell deformation under isotropic pressure and quantified the final cell shape. In this calculation, we evaluated the principal direction of extension for each triangle demonstrated in Figure 6C. With this setting, we captured the local direction of extension in a color diagram (Figure 6D-E, Supplemental Figure S5). We first confirmed that the cell extension anisotropy with vertically longer slits is below 1 (horizontal deformation) regardless of the cell aspect ratio (Figure 6D). Then we tested the cell extension ratio as a function of the slit aspect ratio (Figure 6E), which strikingly showed that slit shape regulates the direction of extension under turgor pressure. The other effects on the cell extension ratio from slit lengths, slit numbers, and slit randomized angle are summarized in Supplemental Figure S5. These simulations revealed that vertically longer slits and their density, size, and parallel alignment enhance horizontal cell extension. These results indicated that the slit structure promotes cell expansion in the direction perpendicular to the slit. Pulvinar slits aligned in the circumferential direction thus have a potential effect on anisotropic cell extension to the PD direction upon turgor pressure, even if anisotropy of the Young’s modulus, like the anisotropy of orientation of cellulose microfibrils, is not assumed.

Finally, we investigated whether pulvinar slits are deformable and how they deform during the change of CMCs in volume. For this experiment, we used the large secondary pulvini from the legume species kudzu (*Pueraria montana var. lobata*). We confirmed that the PD length of CMCs, but not the ML width, is significantly different between hypotonic and hypertonic conditions (Figure 7A), which is consistent with the reported data for other species (Mayer et al., 1985; Koller and Zamski, 2008; Sleboda et al., 2022). In this experiment, we did not detect a negative effect of EGTA treatment (Figure 7A). In addition, we observed pulvinar slits in *P. montana var. lobata* (Figure 7B). We then investigated whether the shape of pulvinar slits is altered when the external solution of slices of CMCs was replaced from a hypotonic solution to a hypertonic solution (hypo/hyper) and vice versa (hyper/hypo). Under the hyper/hypo test condition, CMCs showed PD extension and pulvinar slits appeared to increase their opening width without increasing their size along the ML axis (Figure 7B–C). By contrast, under the hypo/hyper condition, CMCs showed shrinking along the PD axis, and pulvinar slits appeared to decrease their opening width (Figure 7B). These results indicate that changing osmotic pressure makes pulvinar slits reversibly open and close together with the volume change of CMCs.

**Figure 7.**
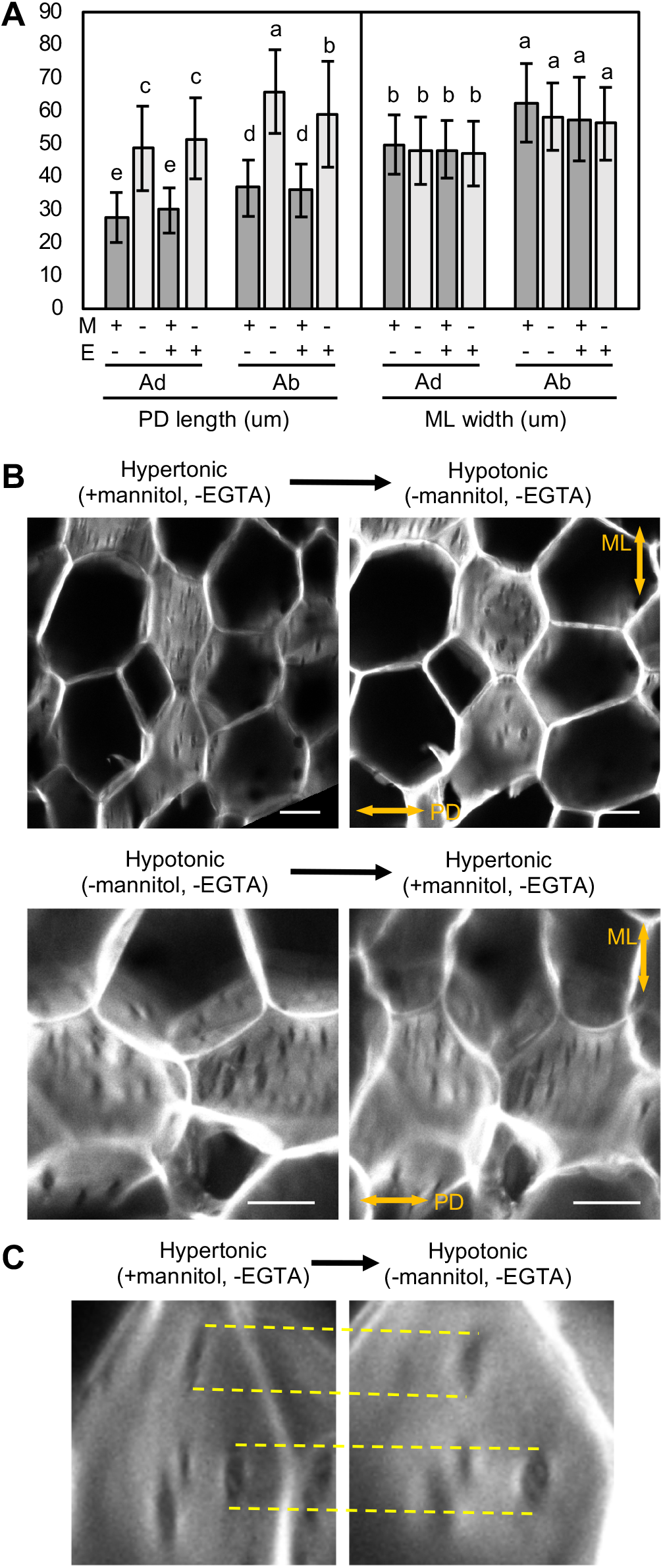
Deformability of pulvinar slits. (A) Size of CMCs in secondary pulvini of *P. montana var. lobata* under hypotonic (− mannitol) and hypertonic condition (+ mannitol). M, mannitol; E, EGTA; Ad, adaxial CMCs; Ab, abaxial CMCs. N = 161 (Ab, −M, −E), 165 (Ab, +M, −E), 242 (Ad, −M, −E), 312 (Ad, +M, −E), 244 (Ad, −M, +E), and 371 (Ad, +M, +E) cells from four leaves, and 207 (Ab, −M, +E) and 315 (Ab, +M, +E) from three leaves. Different letters indicate significant differences (Tukey’s HSD test, family-wise error rate = 0.001). (B) Deformability test of pulvinar slits. The upper panel shows the experiment in response to transfer from hypertonic to hypotonic conditions. The lower panel shows the experiment in response to transfer from hypotonic to hypertonic conditions. (C) Magnified views of the upper panels in B. Yellow dashed lines are in the same positions before and after transfer to the hypotonic condition. Scale bars, 20 μm (B).

## Discussion

The pulvinus is a specialized organ that facilitates dynamic leaf movements. Here, we discovered that CMCs have a unique cell wall structure, designated the pulvinar slit, with a very low content of cellulose or highly methyl-esterified HG while being rich in de-methyl-esterified HG. In addition, we employed computational simulations to predict that the slits bestow CMCs with extensibility in the perpendicular direction. Finally, we demonstrated that pulvinar slits deform in response to changing osmotic pressure. We summarize below the properties of the cell wall and speculate on the roles of pulvinar slits. Finally, we place the discovery of pulvinar slits in context for a greater understanding of plant movement and cell wall mechanics.

We identified pulvinar slits as circumferential slits after calcofluor white staining and showed that the slits are widely conserved in legume species and present in both adaxial and abaxial (i.e., extensor and flexor) CMCs. Whole-mount staining revealed that the structures are not technical artifacts during sectioning but are *bona fide* structures. Staining with pontamine and CBM3a supports to our hypothesis that pulvinar slits reflect the pattern of cellulose deposition. De-methyl-esterified HG, probably without calcium crosslinking, appeared to accumulate abundantly inside pulvinar slits. By contrast, both cellulose and highly methyl-esterified HG were excluded from the pulvinar slits. FTIR analysis and monosaccharide analysis emphasized the distinct cell wall properties of pulvini relative to other axial organs and suggested the abundant deposition of pectin in pulvini. Mayer et al. showed that cellulose microfibrils are oriented circumferentially (Mayer et al., 1985). Observations of pulvinar ultrastructure by transmission electron microscopy identified curving walls and pits including plasmodesmata on CMCs in transverse sections (Fleurat-Lessard and Roblin, 1988; Machado and Rodrigues, 2004; Rodrigues and Machado, 2006). We also observed similar pits here. Vertical sections of pulvini from *G. max* and *M. truncatula* were previously observed by microscopy after staining with toluidine blue O (Gao et al., 2017; Du et al., 2021), which visualizes various components of the cell wall, including pectin (O’Brien et al., 1964; Mitra and Loqué, 2014). However, none of these earlier studies observed the slit-like structures, nor did they report on variation in cell wall thickness. The de-methyl esterification of HG by pectin methylesterases (PMEs) was recently reported to enhance wall hydration and to increase cell wall thickness (Wang et al., 2020). *Medicago truncatula PME12* has been reported to be downregulated in the pulvinus-deficient mutant, *elongated petiolule1* (Bai et al., 2022). Taken together, we propose that de-methyl-esterified HG has a role in thickening the cell wall within the slit, which accumulates little cellulose.

Previous studies using onion epidermal cells suggested that de-methyl esterification of HG by PME treatment causes increased in-plane plasticity of the cell wall but does not affect cell wall creep (Wang et al., 2020). In addition, pectin enhances the strength of the cell wall perpendicular to the plane (Zhang et al., 2019; Wang et al., 2020). However, the addition of calcium blocks the increase of plasticity imparted by PME (Wang et al., 2020). Taken together, we propose that filling the slit with de-methyl-esterified HG without calcium crosslinking achieves both in-plane plasticity and cell wall integrity. In our experiments, we detected a faint signal with the 2F4 antibody (recognizing calcium-crosslinked HG) that did not appear to be associated with pulvinar slits. Although it is reported calcium density in the cell wall can change during leaf movement (Moysset et al., 2019), in our experiment, the calcium chelator EGTA did not have a significant negative effect on the volume of CMCs under both hypotonic and hypertonic conditions. Therefore, calcium has a role in the physiological response to osmotic pressure but may not be involved in mechanical behavior of the cell wall during this deformation. Further analyses will be required to determine the importance of de-methyl-esterified HG inside pulvinar slits and the involvement of the subcellular localization of calcium in the mechanical regulation of CMCs.

Considering the cell wall properties of pulvinar slits, we proposed that the slits are cell wall structures that confer unique mechanical properties, even though a physiological role such as water transport (proposed for cell wall pits by Machado and Rodrigues, 2004; Rodrigues and Machado, 2006) has not yet been excluded. At least within the parameters of Supplemental Table S2, results of FEM suggested that the presence of the slits with low in-plane stiffness can have a potential effect on anisotropic cell extension in the PD direction even if no mechanical anisotropy of the other part, e.g., the anisotropic orientation of cellulose microfibrils, was assumed. The opening width of pulvinar slits increased/decreased under hypo/hypertonic conditions, respectively, indicating the deformability of the slits. Interestingly, pulvinar slits did not appear to develop structural failures at the edges during osmotic changes. This result suggests that the binding between cellulose microfibrils aligned along the ML axis is sufficiently strong at the outside of the edges. The opened slit might have strain energy that causes cell contraction when the difference in osmotic potential between the inside and the outside of the CMC decreases. In other words, the opening and closing of pulvinar slits might be the driving force for the elastic behavior of the CMC. Taken together, we provide a hypothesis that pulvinar slits have a role in dynamic leaf movement through repetitive and reversible deformation of CMCs in concert with other factors, including cellulose orientation (Mayer et al., 1985), pectin-rich composition of the cell wall (this study), the geometry of CMCs, and the actin cytoskeleton (Kameyama et al., 2000; Kanzawa et al., 2006). There is no direct evidence to indicate the extent to which pulvinar slits contribute to the elastic behavior of the CMCs, so caution is warranted in our interpretation. Nevertheless, the wide conservation of pulvinar slits in legume species suggests that they are at least advantageous for plant survival. Since the primary function of pulvini is leaf movement, it is reasonable to hypothesize that pulvinar slits improve plant fitness by contributing to leaf movement. The mechanical contribution of pulvinar slits should be verified by genetic analysis in the future.

In recent years, the cellular basis of repeated and reversible cell deformation driven by turgor has been extensively studied using stomata. Stomata are composed of a pair of guard cells on the leaf epidermis that control gas exchange and repeatedly adjust their aperture by modulating the activity of transmembrane ion channels in response to environmental signals (Kollist et al., 2014). Guard cells have unique morphological features, for example, a kidney-like shape in many eudicots or a dumbbell-like shape in grasses. Major components of primary cell walls, e.g., cellulose, xyloglucan, and pectin, contribute to stomatal movements (Rui and Anderson, 2016; Amsbury et al., 2016; Yi et al., 2018), and computational model–based analyses have demonstrated that cellulose-based anisotropy and polar stiffening of stomata are important factors (Carter et al., 2017; Woolfenden et al., 2017; Woolfenden et al., 2018). Notably, de-methyl esterification of HG is required for stomatal function (Amsbury et al., 2016). The cell wall of guard cells is low in highly methyl-esterified HG and rich in de-esterified HG, a composition similar to that of the inside of pulvinar slits. Importantly, the cell shape, the spatial arrangement of cell wall components within a cell, and the homogeneity of cell wall thickness are significantly different between CMCs and guard cells. These differences may be closely related to those seen in the directions of cell deformation: anisotropic extension of CMCs to the PD direction and opening/closing of guard cells.

Intriguingly, one of the characteristics of slits of the PCW in pulvinar CMCs, e.g., scattered regions with less cellulose, is similar to that of pits of the secondary cell wall in xylem vessels (Oda and Fukuda, 2012a). The deposition pattern of the secondary cell wall is controlled through the localization of cellulose synthase complexes by MICROTUBULE DEPLETION DOMAIN 1 and the Rho-GTPase ROP in metaxylem cells (Oda and Fukuda, 2012b). By analogy, the unique cell wall pattern seen in the CMCs of pulvini might be controlled by the localization of cellulose synthase complexes as well. However, unlike xylem vessels with a thicker secondary cell wall, variation in cell wall thickness has not been reported in the CMCs of pulvini. Furthermore, the xylem pits have xylan deposits (Wang et al, 2022), which is different from pulvinar slits depositing de-methyl-esterified HG inside. These differences suggest that the underlying mechanism establishing the unique cell wall pattern of CMCs may be considerably different from that of xylem vessels.

Furthermore, from a broader perspective, it is intriguing that CMCs with pulvinar slits are structurally analogous to kirigami (the Japanese art of paper cutting), an example of which has many straight cuttings, i.e., slits. Slits of kirigami increase extensibility of the sheet and prevent unpredictable local structural failures (Shyu et al., 2015; Isobe and Okumura, 2016). In conclusion, we propose that pulvinar slits are an example of specialization in motor organs. Our findings illustrate the structural diversity of the cell wall and its association with the mechanical regulation of plant movement.

## Methods

### Plant materials

Plants used in this study were grown under four different conditions: outdoor collection of wild plants, outdoor cultivation, indoor cultivation, and greenhouse cultivation (Supplemental Table S1). Wild plants of *Desmodium paniculatum*, Persian silk tree (*Albizia julibrissin*), white clover (*Trifolium repens*), and kudzu (*Pueraria montana var. lobata*) were collected outdoors in Nara, Japan (34°43.5□N, 135°44.1□E) in the summer of 2019, 2021, and 2022. Adzuki bean (*Vigna angularis*) Dainagon (TOHOKU SEED Co., Ltd, Japan), cowpea (*Vigna unguiculata*) Sansyaku (TOHOKU SEED Co., Ltd, Japan), common bean (*Phaseolus vulgaris*) L. cv Morocco (Atariya Noen Co. Ltd, Japan or TAKII & Co., Ltd, Japan), and sword bean (*Canavalia gladiata*) (Atariya Noen Co. Ltd, Japan or TAKII & Co., Ltd, Japan) were grown outdoors and collected in the summer of 2019. Sensitive plant (*Mimosa pudica*) Wase (SAKATA SEED CORPORATION, Japan), common licorice (*Glycyrrhiza glabra*) (purchased at a garden store), and carob (*Ceratonia siliqua*) (purchased at a garden store) were grown under white LED light (SMD 5050; Samsung) at 100–200 μmol m^−2^ s^−1^ under long-day conditions (16 h light/8 h dark). Plants of soybean (*Glycine max*) and birdsfoot trefoil (*Lotus japonicus*) were grown in the greenhouse at NAIST under a photoperiod adjusted to long-day conditions by supplementary light. Except for species collected outdoors, potted plants of all tested species were grown on culture soil (Kohnan Shoji Co., Ltd. or Nihon Hiryo Co., Ltd). Samples were harvested at Zeitgeber time (ZT) 2–7 (dawn being ZT0).

### Staining and microscopy

Leaves with pulvini were fixed in FAA solution (5% [v/v] formaldehyde, 5% [v/v] acetic acid, 45% [v/v] ethanol) or PFA solution (2.22% [w/v] paraformaldehyde, 0.5 M mannitol, 50 mM PIPES, pH 6.9) and stored at 4°C. For the analysis of *D. paniculatum* (Figure 1) and nine other species (excluding *L. japonicus*), fixed samples were transferred into distilled water several hours before analysis and sliced with a single-edged razor. For analyses of *D. paniculatum* (Figures 2, 3, Supplemental Figure S1) and *L. japonicus*, fixed samples were washed in distilled water several times, embedded in 5% (w/v) agarose gel, and sliced into 100-μm sections with a MicroSlicer Zero 1 (DOSAKA EM, Kyoto, Japan).

For calcofluor white staining, the sections were placed onto glass slides, treated with one drop of calcofluor white stain solution (Sigma-Aldrich, USA) and one drop of 10% (w/v) potassium hydroxide for 1 min, mounted in distilled water, and covered by a cover slip, before observations with a confocal laser-scanning microscope (LSM800, Zeiss, Germany) equipped with a Plan-Apochromat 20×/0.8 M27 objective lens (Zeiss, Germany; excitation 405 nm; emission 400–560 nm). For whole-mount observations of *L. japonicus* pulvini, fixed samples were transferred into 1× phosphate-buffered saline (PBS), pH 7.5 (130 mM NaCl, 7 mM Na_2_HPO_4_, and 3 mM NaH_2_PO_4_), washed in 1× PBS twice, soaked in ClearSee solution (10% [w/v] xylitol, 15% [w/v] sodium deoxycholate, and 25% [w/v] urea) (Kurihara et al., 2015) for 1 week, stained with 2.5 μg/mL calcofluor white in ClearSee solution for 5 d, mounted in ClearSee solution, and observed under a LSM800 microscope as described above.

For pontamine staining, the sections were incubated with 50 mg/L Direct Fast Scarlet 4BS (Wako, Japan) for 20 min, rinsed with 100 mM NaPO_4_ buffer several times, mounted in distilled water, and covered by a cover slip, before observations with a LSM800 microscope equipped with a Plan-Apochromat 20×/0.8 M27 objective lens (excitation 488 nm; emission 560–605 nm).

For immunostaining, sections were rinsed with Tris-buffered saline (TBS, 20 mM Tris-HCl, 150 mM NaCl, pH 7.5) three times, incubated in blocking solution (1% [w/v] bovine serum albumin, 1% [w/v] normal goat serum (Abcam ab7481) in TBS with 0.2% [v/v] Tween-20 [TBST]), rinsed with TBST three times, treated with TBS (for no antibody control) and with primary antibodies (CMB3a, LM15, LM19, LM20, 2F4, PlantProbes, diluted 1:10) in TBS at 4°C for 1 h, rinsed with TBST three times, incubated with secondary antibodies (anti-rat IgG-CFL488, anti-mouse IgG-Alexa488, anti-His tag IgG-CFL488, all diluted 1:100) at 4°C for 1 h, and rinsed with TBST five times. The samples were mounted in 1 mL TBST with 10 μL calcofluor white stain solution and 10 μL of 10% (w/v) potassium hydroxide, covered by a cover slip, and observed with a LSM800 microscope equipped with a Plan-Apochromat 20×/0.8 M27 objective lens (excitation 488 nm; emission 400–575 nm; excitation 405 nm; emission 400–510 nm).

To test the deformability of pulvinar slits, tissue slices of CMCs were excised from the abaxial side of fresh pulvini of *P. montana var. lobata*, immediately transferred to 1 mL of 10 mM PIPES-KOH, pH 7.0 (stabilizing buffer [SB]), with or without 450 mM mannitol after adding 10 μL of calcofluor white stain solution, and immersed for 20 min or more. The stained samples were placed in a glass-bottom dish, and calcofluor-while fluorescence was observed as with fixed samples. Then, 30 μL of the SB solution alone or with 450 mM mannitol (made fresh) was applied to the samples, and after a 20-min incubation, the same positions on the slices were observed.

### Cell wall analysis

Primary pulvini, secondary pulvini, petioles, young stems, and mature stems of *D. paniculatum* were harvested for analysis of cell wall composition. Preparations of cell wall residues and monosaccharide composition analysis were performed as previously described (Sakamoto et al., 2015). Fourier-transform infrared spectroscopy (FTIR) analysis was performed as described previously (Sakamoto et al., 2018).

### Compression test of pulvini and other axial organs

Each organ was excised into segments of approximately 2 mm in length just before the test. Load while under a constant-speed deformation (0.05 mm/s) was measured with a RHEONERII creep meter (Yamaden, Japan; RE2-3305C) equipped with a 20 N load cell. Excised specimens were placed on a moving platform with the apex-base axis as the horizontal axis and compressed by sandwiching each specimen between the moving platform and a cylindrical plunger with a diameter of 3 mm. Strain was calculated by dividing the deformation by the height at time 0. Analyses were conducted in the daytime on the same day as sample collection.

To measure PD extension, videos were captured during the compression test with a 10–200 × magnification USB digital microscope (Giwox, China) and the PhotoBooth app (Apple, USA) or a HiVision Microscope FZ200BA equipped with HDMI recorder X Capture MINI (Shodensha, Japan). Captured videos were divided into frames with ffmpeg software (https://ffmpeg.org/). Images at the time of each 10% strain were selected based on their frame rate and the height at time 0. The PD length of each organ was manually measured with ImageJ (Fiji; https://fiji.sc/), and the extension ratio was calculated by the following equation in Microsoft Excel.

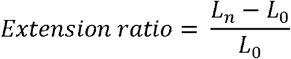

where *L*_0_ represents the length when strain is 0, and *L_n_* represents the length when strain is *n* (0.1–0.8).

For observations of loaded pulvini (Supplemental Figure S4), prior to observation, tissues were manually compressed with a 10 N load while measuring with an electronic balance. For observations by scanning electron microscopy, tissues were frozen in liquid nitrogen for 15 s and were observed with a FEI Quanta 250 scanning electron microscope (ThermoFisher Scientific, USA). For observations by confocal laser-scanning microscopy, fresh samples embedded in 6% (w/v) agarose gel were sliced into 50-μm sections with a MicroSlicer Zero 1. Sections were observed with a LSM800 microscope.

### Computational modeling

A continuous mechanical model for the CMCs was constructed based on the finite-element method (Bonazzi et al., 2014; Hong et al., 2016; Tsugawa, 2020). Details are provided in Supplemental material. We considered different initial cell shapes as well as different slit shapes, sizes, and orientations. Cells were examined with different initial shapes close to the regular hexagon as defined above with four vertical slits (Figure 6D). Cells with different aspect ratios of slits (Figure 6E) and with different numbers of slits, slit areas, and randomized angles of slits were also examined (Supplemental Figure S5). The code used for modeling is available at GitHub (https://github.com/SatoruTsugawa/hexagon-model).

### Statistical analyses

Microsoft Excel for Mac v16.16.11 (Microsoft, USA), R (https://www/r-project.org), or the Python language v3.7.2 (https://www.python.org/) and Python packages/libraries including Pandas v1.1.3 (https://pandas.pydata.org/), NumPy v1.19.4 (https://www.numpy.org/), Matplotlib v3.3.2 (https://matplotlib.org/), Seaborn v0.11.0 (https://seaborn.pydata.org/), Scikit-learn v0.23.2 (https://scikit-learn.org/), and Scikit-posthocs v0.6.6 (https://github.com/maximtrp/scikit-posthocs) were used for statistical analyses. The Tukey’s HSD test was performed after the ANOVA test. The details are given in Supplemental Table S3.

## Abbreviations

CMC: cortical motor cell
FTIR: Fourier-transform infrared spectroscopy
PCW: primary cell wall
SCW: secondary cell wall
HG: homogalacturonan
PD: proximo-distal
CBM3a: carbohydrate-binding module 3
FEM: finite-element method
PME: pectin methylesterase

## Acknowledgements and funding

We thank Kiyotaka Okada (Ryukoku Univ., Japan) and Nobutaka Mitsuda (AIST, Japan) for helpful comments, Atsunobu Suzuki (NAIST, Japan), Hitomi Ichikawa (NAIST, Japan), Hiroyo Aoki (NAIST, Japan) and Aeni Hosaka (AIST, Japan) for technical support. We thank Tadashi Kunieda (NAIST, Japan), Ko Kato (NAIST, Japan) and members of the Demura lab for general support during experiments. *G. max* seeds were obtained from LegumeBase (http://www.legumebase.brc.miyazaki-u.ac.jp/), which is supported by the National BioResource Project (NBRP) in Japan.

This work was supported by JSPS KAKENHI Grant Numbers JP20K15832 (to S.T.), JP19K16174 (to S.S.), JP19K23753 and JP20K06707 (to M.N.), the Japan Science and Technology Agency [CREST(JPMJCR2121)] (to S.T.), and by MEXT KAKENHI Grant-in-Aid for Scientific Research on Innovative Areas “Plant-Structure Optimization Strategy” Grant Numbers JP18H05484 and JP18H05489 (to T.D.).

## Author contributions

M.T. and M.T.N. contributed the conception and design of the study, performed biological/histochemical/mechanical experiments and image/statistical analyses, and wrote the first draft of the manuscript. S.T. performed FEM analysis and wrote the section on FEM simulations. S.S. performed FTIR and monosaccharide analyses, contributed to data interpretation, and wrote the section on the cell-wall assays. T.D. contributed significantly to the revision of the experimental design and the revision of an earlier version of the manuscript. All authors discussed the results and contributed to the final manuscript.

## Supplemental information

Supplemental figures, tables and material.

