## Supplemental material for "Pulvinar Slits: Cellulose-deficient and De-Methyl-Esterified Homogalacturonan-Rich Structures in a Legume Motor Cell"

**Computational modeling**

We constructed a continuous mechanical model for the CMCs based on the finite-element method (Bonazzi et al., 2014; Hong et al., 2016; Tsugawa, 2020). Only the cross section of cells was modeled, whereby a two-dimensional medium with a prescribed distribution of elastic modulus $E$ and Poisson’s ratio $\nu$ is deformed. In the real situation of a cell, the cell medium is surrounded by the cell wall, but we only focused here on the deformation of the medium inside the cell wall. Cell deformation occurs by increments in the cell area: The initial resting shape is inflated by turgor pressure $P$, leading to a new shape at equilibrium. The model was implemented and solved with solvers in FreeFem++ (Hecht, 2019), and the results were analyzed using gnuplot scripts. There were on average about 2500 triangular elements per cell in our model. This fine distribution of triangles allowed us to avoid some abnormal deformations at the edges or at the corners. In this study, cell deformation with some inner structure of the cell under uniform pressure was examined, which indicated a mechanically unstable orientation depending on the internal structure of the cell. We also performed a computational compression test of the CMCs at the cellular level.

We used the generalized Hooke’s law with the stress tensor $\sigma$ and the strain tensor $\varepsilon$ through the elasticity matrix:

$$\left( \begin{matrix} \sigma_{xx} \\ \sigma_{yy} \\ \sigma_{xy} \end{matrix} \right)=\left( \begin{matrix} A & B & 0 \\ B & A & 0 \\ 0 & 0 & C \end{matrix} \right)\left( \begin{matrix} \varepsilon_{xx} \\ \varepsilon_{yy} \\ \varepsilon_{xy} \end{matrix} \right)$$

where $A=\frac{1}{1-\nu^{2}}E, B=\frac{\nu}{1-\nu^{2}}E,C=\frac{1}{1+\nu}E$ ($\sigma_{\mathrm{zz}}=0$, longitudinal deformation in Section 13 of Landau and Lifshitz, 1986). The starting configuration was a regular hexagon or some similar shape. The side of the hexagon was set to 1 arbitrary unit (a.u.), which corresponds to the actual cell side of the 20-μm scale bars in Figure 1. Young’s modulus of CMCs was assumed as $E=1$ MPa and Poisson’s ratio as $\nu=0.35$ based on a previous study in flexor tissue ($E=0.6\sim1.4$MPa; Mayer et al., 1985) and Arabidopsis sepals ($E=3.27$ MPa and $\nu=0.48$; Hong et al., 2016). In practice, we first simulated the deformed shape for all combinations of the parameters $E=1, 5, 10$ (MPa) and $\nu=0.35, 0.40, 0.45$ and confirmed that all cases were qualitatively similar. Turgor pressure of CMCs was assumed to be $P=0.1$ (MPa), which is a slightly lower value but on the same order of the previously reported pressure value 0.3~1.0 MPa (Cosgrove, 1993; Lintilhac et al., 2000; Radotić et al., 2012). With this value, we mimicked a slight change of turgor pressure. We confirmed that the deformed shape for the parameters $P=0.1\sim1.0$ (MPa) was qualitatively similar.

The cell geometry was defined as a regular hexagon with side = 1 (a.u.) with the origin as the center of gravity of the cell. We also considered similar cell shapes to the regular hexagon in which the coordinates $(x,y)$ are multiplied by $(\alpha x,y)$, where $\alpha$ is the cell aspect ratio (Figure 6D). The slit geometry was defined as an ellipse with the coordinates $(x,y)$ satisfying $\frac{\left( x-x_{s} \right)^{2}}{r_{x}^{2}}+\frac{\left( y-y_{s} \right)^{2}}{r_{y}^{2}}=1,$ where $(x_{s},y_{s})$ is the origin of the slit, and $r_{x}$ and $r_{y}$ are the half of major width and the half of minor width, respectively. We located the i-th origin of the slits as $\left( x_{s}^{i},y_{s}^{i} \right)=\left( \cos\left( \frac{2\pi(i-1)}{N} \right),\sin\left( \frac{2\pi(i-1)}{N} \right) \right), i=1,\cdots,N, N\in2\mathbb{N.}$ We assumed that the elastic properties inside the slits are negligibly small, which allowed us to make the mesh in the slits empty (i.e., E = 0 MPa).

**Bonazzi D, Julien JD, Romao M, Seddiki R, Piel M, Boudaoud A, Minc N** (2014) Symmetry breaking in spore germination relies on an interplay between polar cap stability and spore wall mechanics. Dev Cell **28:** 534-546

**Cosgrove DJ** (1993) Wall extensibility: its nature, measurement and relationship to plant cell growth. New Phytol **124(1):** 1-23

**Hecht F** (2019) Freefem++ (Third Edition, Version 3.54 http://www.freefem.org/ff++/)

**Hong L, Dumond M, Tsugawa S, Sapala A, Routier-Kierzkowska AL, Zhou Y, Chen C, Kiss A, Zhu M, Hamant O, Smith RS, Komatsuzaki T, Li CB, Boudaoud A, Roeder AH** (2016) Variable Cell Growth Yields Reproducible Organ Development through Spatiotemporal Averaging. Dev Cell **38:** 15-32

**Landau LD, Lifshitz E** (1986) Theory of Elasticity. Pergamon Press, Oxford.

**Lintilhac PM, Wei C, Tanguay JJ, Outwater JO** (2000) Ball Tonometry: A Rapid, Nondestructive Method for Measuring Cell Turgor Pressure in Thin-Walled Plant Cells. J Plant Growth Reg **19:** 90–97

**Mayer WE, Flach D, Raju MV, Starrach N, Wiech E** (1985) Mechanics of circadian pulvini movements in *Phaseolus coccineus* L.: Shape and arrangement of motor cells, micellation of motor cell walls, and bulk moduli of extensibility ([Formula: see text]). Planta **163:** 381-390

**Radotić K, Roduit C, Simonović J, Hornitschek P, Fankhauser C, Mutavdžić D, Steinbach G, Dietler G, Kasas S** (2012) Atomic Force Microscopy Stiffness Tomography on Living Arabidopsis thaliana Cells Reveals the Mechanical Properties of Surface and Deep Cell-Wall Layers during Growth. Biophys J **103(3):** 386–394

**Tsugawa S** (2020) Suppression of soft spots and excited modes in the shape deformation model with spatio-temporal growth noise. J Theor Biol **486:** 110092
