## Supplemental Figures for "Pulvinar Slits: Cellulose-deficient and De-Methyl-Esterified Homogalacturonan-Rich Structures in a Legume Motor Cell"

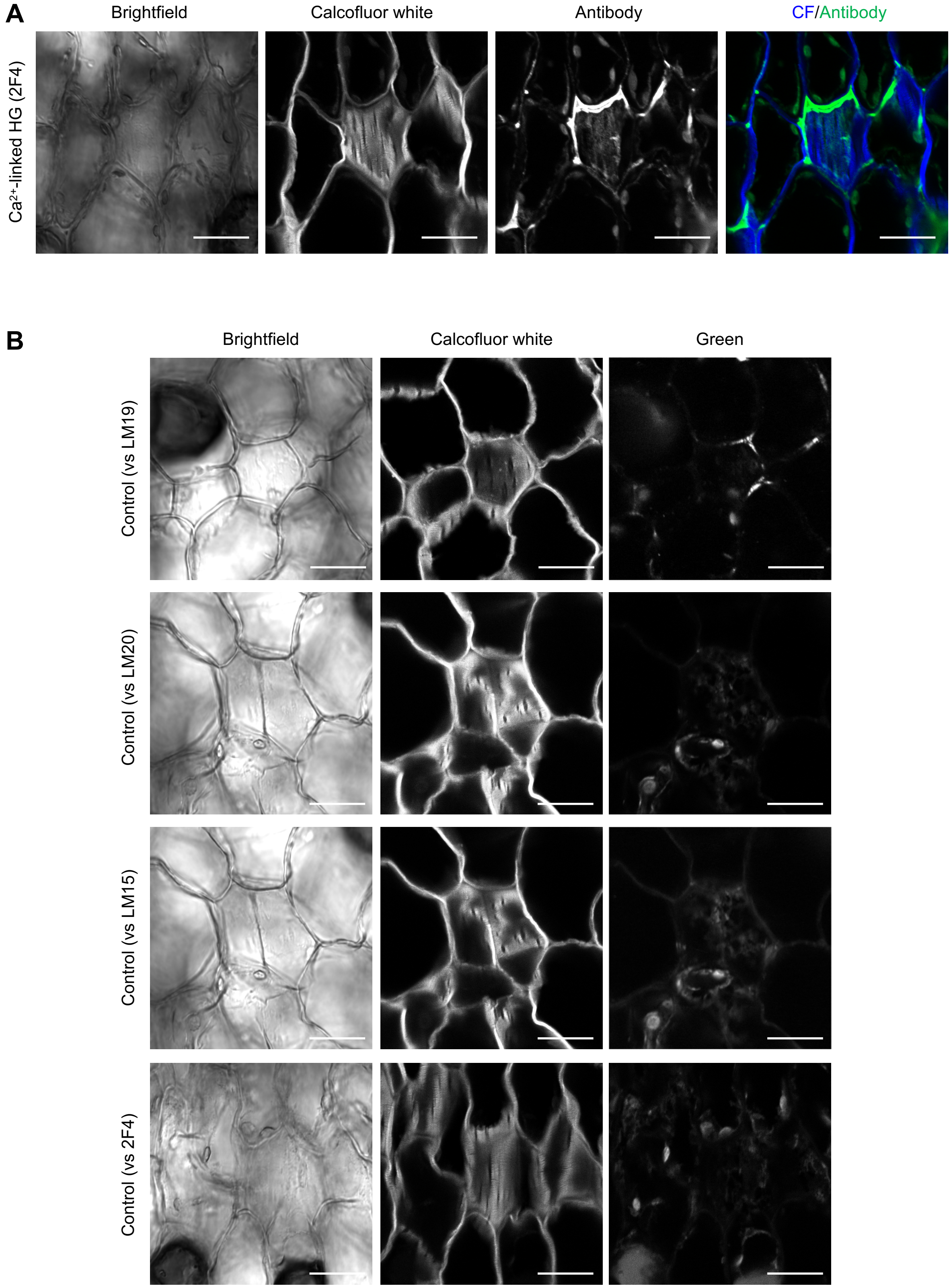


**Supplemental Figure S1**

(A) The signal of the 2F4 antibody in CMCs. (B) Negative controls treated only with secondary antibodies. Each control was observed under the same imaging conditions as the corresponding primary antibody treatment conditions. Images of negative control for the LM15 and LM20 antibodies are derived from the same region but the imaging conditions correspond to those for each antibody. Scale bars, 20 $\mu$m.

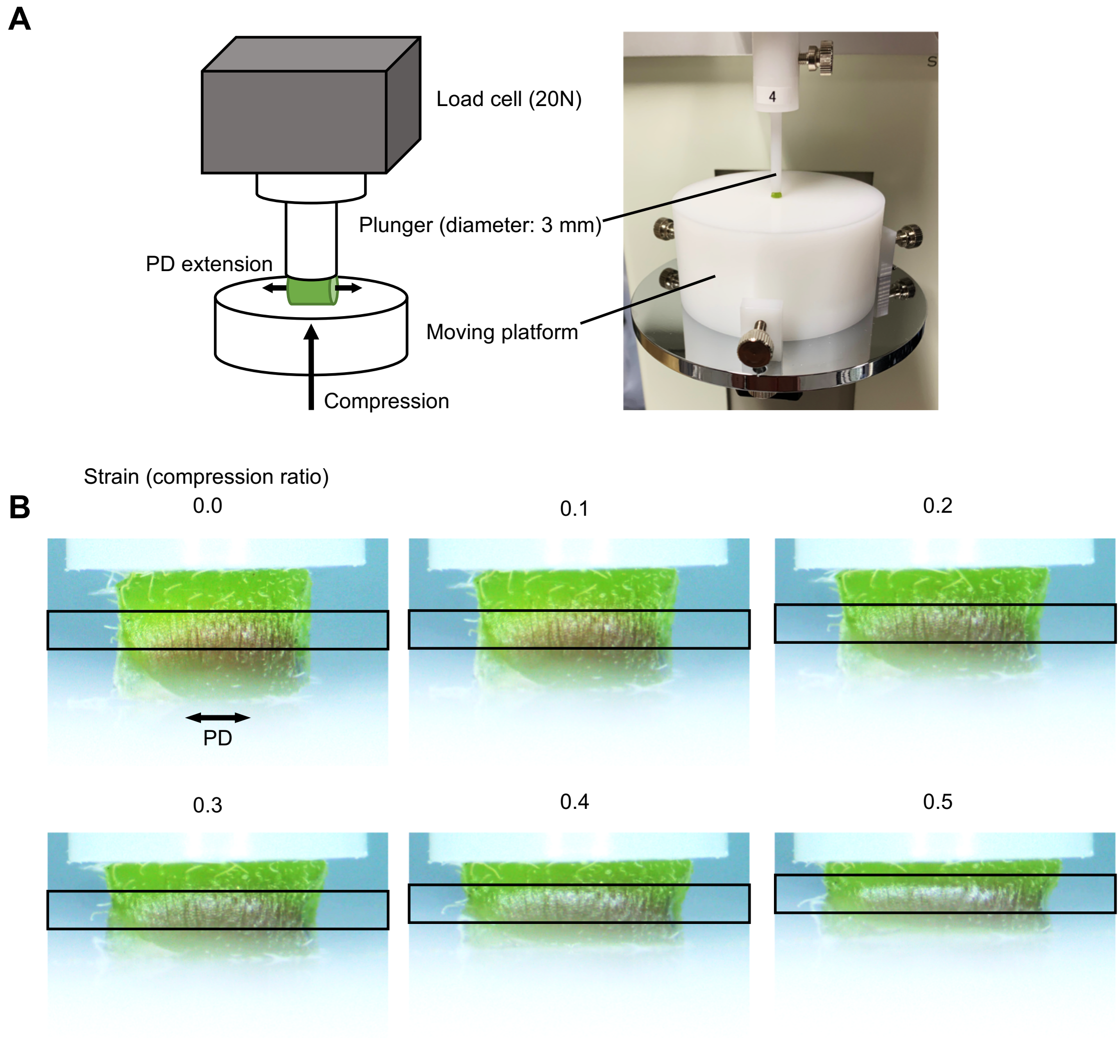


**Supplemental Figure S2**

(A) Schematic diagram and a raw image of compression test. (B) Whole images of Figure 6D. Black boxes show the cut out part. PD, the proximo-distal axis.

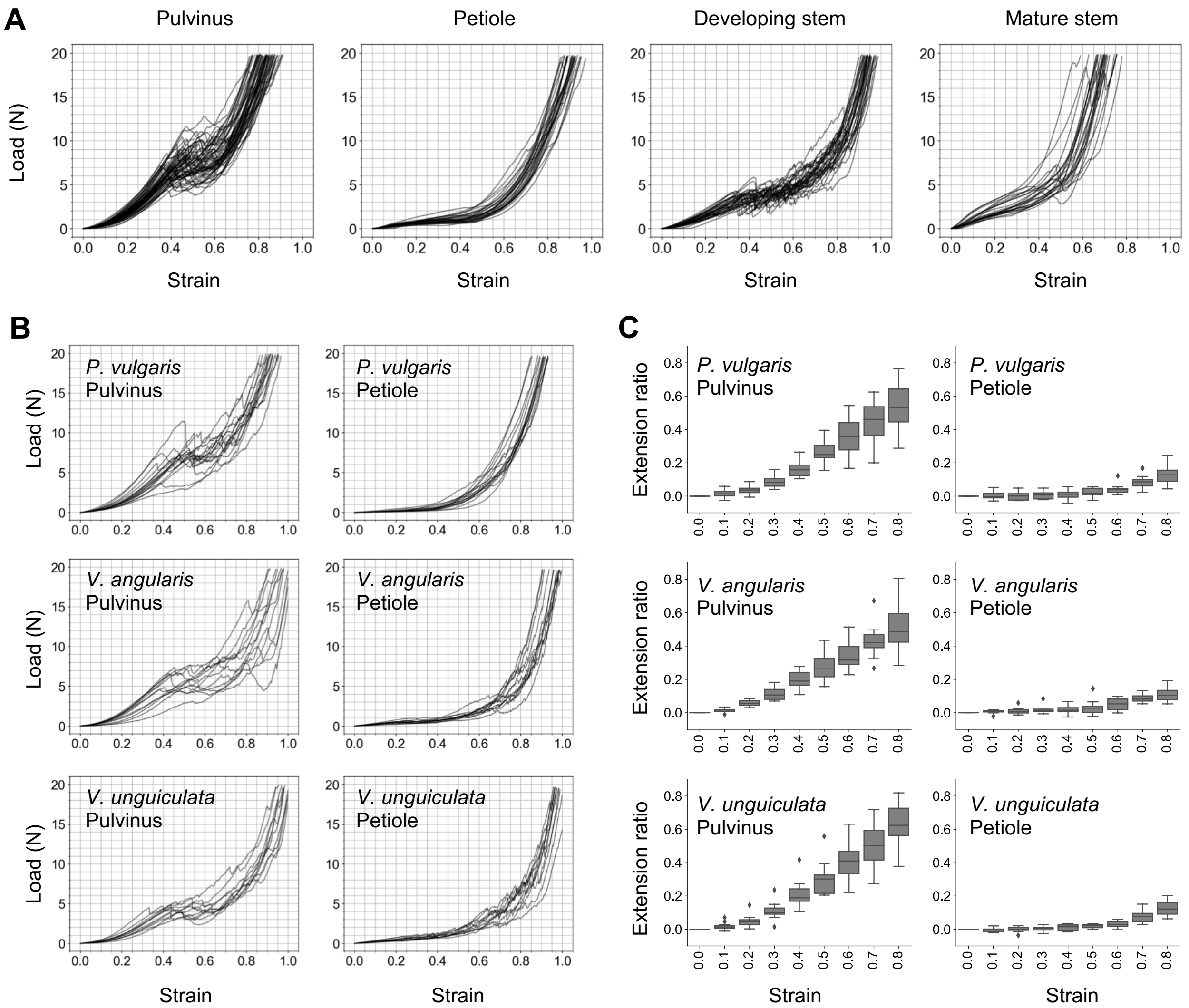


### Supplemental Figure S3

(A) The load-strain curves when compressing each organ of *D. paniculatum*. N = 60 (pulvini), 29 (petioles), 33 (developing stems) and 24 (mature stems). (B) The load-strain curves when compressing pulvinus and petioles. (C) The proximo-distal extensibility of pulvinus and petioles. In B and C, N = 15 (petioles of *P. vulgaris*), 16 (pulvini of *P. vulgaris*), 12 (*V. angularis*) and 11 (*V. unguiculata*).

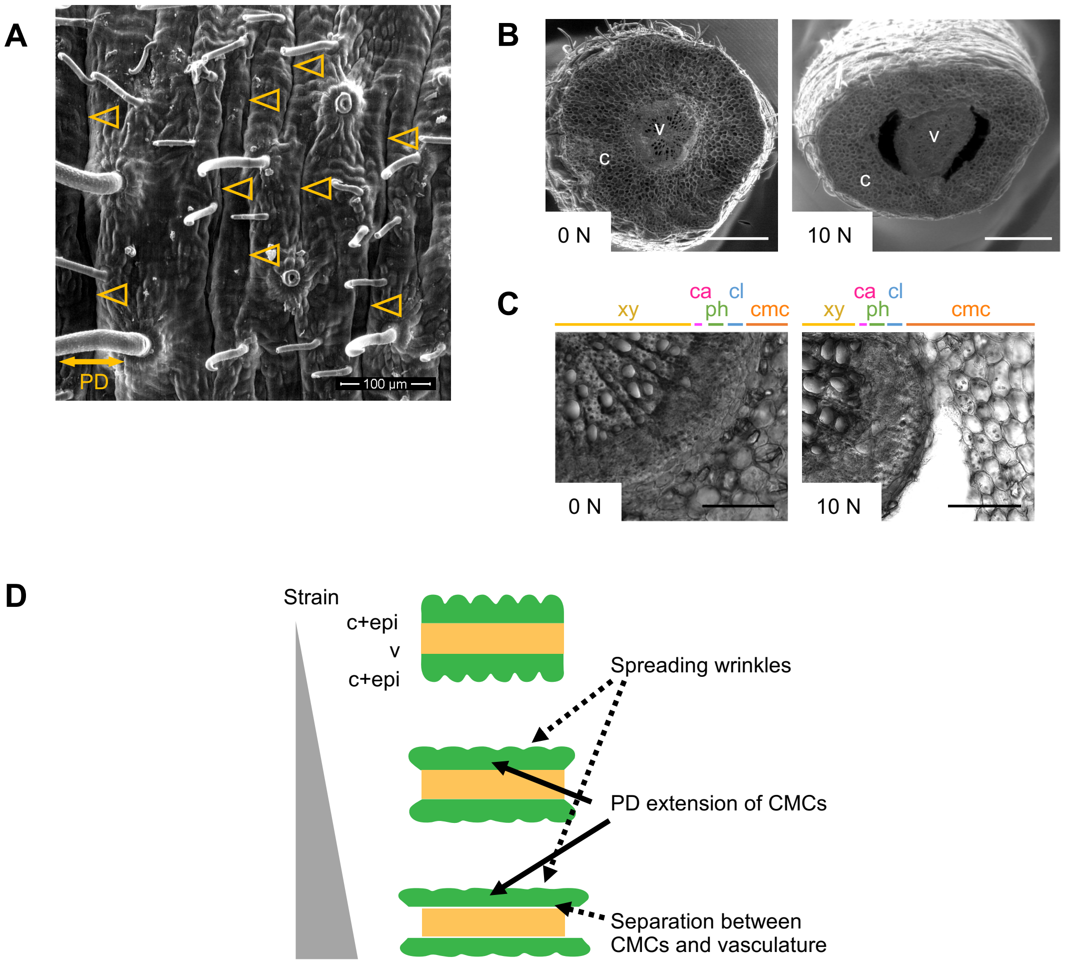


**Supplemental Figure S4**

(A) A scanning electron micrograph of the surface of a *D. paniculatum* pulvinus. Arrowheads represent tiny wrinkles that aligned perpendicularly to the PD axis. (B) Scanning electron micrographs of pulvini without compression (0 N) or after compression (10 N). v, vasculature; c, cortex. (C) Cross sections of *D. paniculatum* pulvini without compression (0N) or after compression (10N) observed by confocal laser scanning microscopy. Xy, xylem; ca, cambium; ph, phloem; cl, collenchyma; cmc, cortical motor cells. (D) A model of the events that occur in compression test. The illustrations indicate the longitudinal section of a pulvinus. c+epi, CMCs and epidermis; v, vasculature. Scale bars, 500 $\mu$m (B), 100 $\mu$m (C).

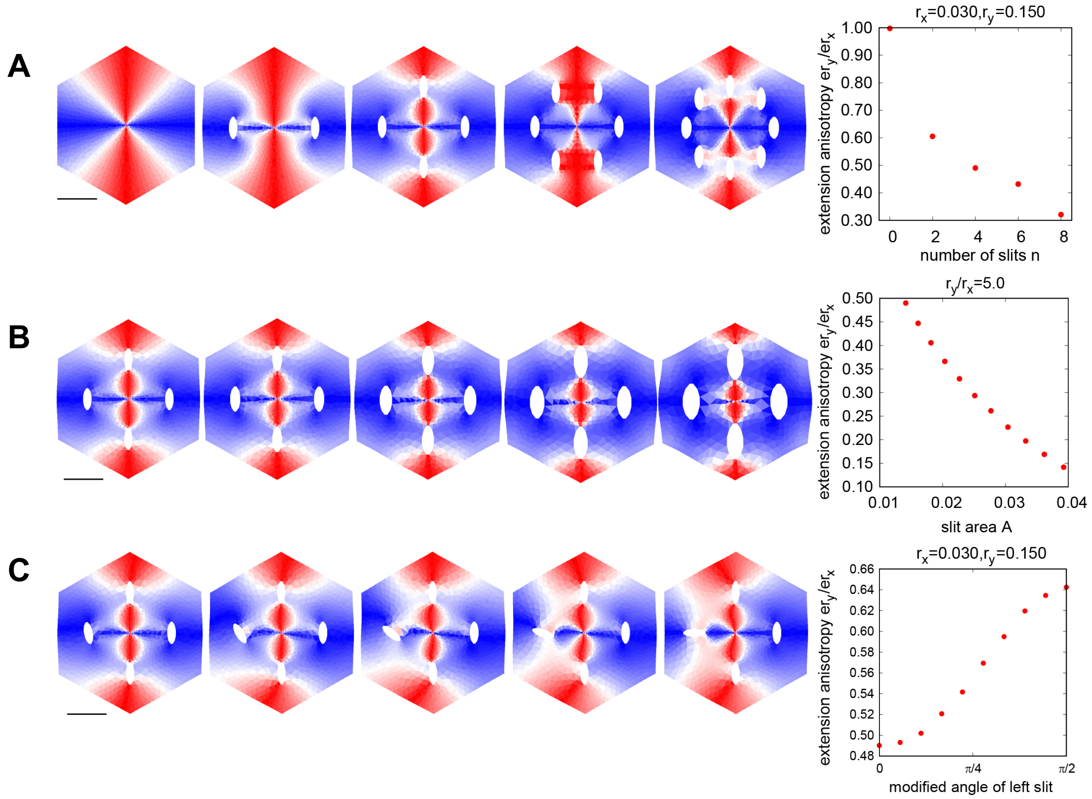


### Supplemental Figure S5

(A-C) Results of FEM simulations to clarify the effect of slit number (A), slit size (B), or slit parallelism (C). Color label of left panels is shown in Figure 6C.
