## Supplemental Tables for "Pulvinar Slits: Cellulose-deficient and De-Methyl-Esterified Homogalacturonan-Rich Structures in a Legume Motor Cell"

Supplemental Table S1. Taxon of tested species

| Family | Subfamily | Subclade | Tribe | Species | Preparation |
|---|---|---|---|---|---|
| Fabaceae | Caesalpinioideae | | Caesalpinieae | *Ceratonia siliqua* | a |
| | | Mimosoid | Mimosoideae | *Mimosa pudica* | a |
| | | | | *Albizia julibrissin* | b |
| | Papilionoideae | Indigoferoid/Millettioid | Desmodieae | *Desmodium paniculatum* | b |
| | | | Diocleae | *Canavalia gladiata* | c |
| | | | Phaseoleae | *Glycine max* | d |
| | | | | *Phaseolus vulgaris* | c |
| | | | | *Vigna angularis* | c |
| | | | | *Vigna unguiculata* | c |
| | | | | *Pueraria montana var. lobata* | b |
| | | Robinioid | Loteae | *Lotus japonicus* | d |
| | | IRLC | Galegeae | *Glycyrrhiza glabra* | a |
| | | | Trifolieae | *Trifolium repens* | b |

a, indoor cultivation; b, outdoor collection of wild plants; c, outdoor cultivation; d, greenhouse cultivation.

Supplemental Table S2. Parameters for FEM simulation

| Parameter | Tested range |
|---|---|
| Elastic modulus | 1, 5, 10 MPa |
| Poisson's ratio | 0.35, 0.40, 0.45 |
| Turgor pressure | 0.1-1.0 MPa |
| Cell aspect ratio ($L_y/L_x$) | 0.4-1.8 |
| Slit aspect ratio ($r_y/r_x$) | 0-10 |
| Number of slits | 0-8 |
| Slit area | 0.01-0.04 |
| Modified angle of left slit | 0-π/2 |

Supplemental Table S3. Computational tools for each statistical analysis

| Analysis | | Tool | Function |
|---|---|---|---|
| 95% confidence interval | Figure 4A | Python Seaborn | lineplot |
| Principal component analysis (PCA) | Figure 4B | Python Scikit-learn | decomposition.PCA |
| Heatmap with dendrogram | Figure 4C | Python Seaborn | clustermap |
| Kruskal-Wallis H-test | Table 1 | Python Scipy | stats.kruskal |
| Dunn’s test (Holm's method) | Table 1 | Python Scikit-posthocs | posthoc_dunn(p_adjust = 'holm') |
| Tukey's HSD test (and ANOVA) | 5A-B, 7A | R | TukeyHSD, aov |
| Two-sided Welch’s t-test | 5C | Python Scipy | stats.ttest_ind |
